# Autism excitation-inhibition imbalance linked to brain hyperconnectivity: An analysis based on 657 autistic subjects

**DOI:** 10.1101/2022.07.14.500131

**Authors:** Javier Rasero, Antonio Jimenez-Marin, Ibai Diez, Roberto Toro, Mazahir T. Hasan, Jesus M. Cortes

**Affiliations:** Cognitive Axon Lab, Department of Psychology, Carnegie Mellon University, Pittsburgh, United States of America; Computational Neuroimaging Lab, Biocruces-Bizkaia Health Research Institute, Barakaldo, Spain; Biomedical Research Doctorate Program, University of the Basque Country (UPV/EHU), Leioa, Spain; Department of Radiology, Division of Nuclear Medicine and Molecular Imaging, Massachusetts General Hospital and Harvard Medical School, Boston, United States of America; Gordon Center for Medical Imaging, Department of Radiology, Massachusetts General Hospital and Harvard Medical School, Boston, United States of America; Athinoula A. Martinos Center for Biomedical Imaging, Massachusetts General Hospital, Harvard Medical School, Boston, United States of America; Institut Pasteur, Université de Paris, Département de neuroscience, F-75015 Paris, France; Laboratory of Brain Circuits Therapeutics, Achucarro Basque Center for Neuroscience, Leioa, Spain; IKERBASQUE, The Basque Foundation for Science, Bilbao, Spain; Department of Cell Biology and Histology, University of the Basque Country (UPV/EHU), Leioa, Spain

**Keywords:** Autism, Subtyping, Excitation-Inhibition Balance, Hyperconnectivity, Allen Human Brain Atlas

## Abstract

The large heterogeneity in autism spectrum disorder (ASD) is a major drawback for the development of therapies. Here, we apply consensus-subtyping strategies based on functional connectivity patterns to a population of N=657 quality-assured autistic subjects. We found two major subtypes (each divided hierarchically into several minor subtypes): Subtype 1 exhibited hypoconnectivity (less average connectivity than typically developing controls) and subtype 2, hyperconnectivity. The two subtypes did not differ in structural imaging metrics in any of the regions analyzed (64 cortical and 14 subcortical), nor in any of the behavioral scores (including Intelligence Quotient, ADI and ADOS). Finally, we used the Allen Human Brain Atlas of gene transcription to show that subtype 2, corresponding with about 42% of all patients, had significant enrichment (after multiple comparisons correction) to excitation-inhibition (E/I) imbalance, a leading reported mechanism in the developmental pathophysiology of ASD. Altogether, our results support a link between E/I imbalance and brain hyperconnectivity in ASD, an association that does not exist in hypoconnected autistic subjects.

## Introduction

Autism encompasses multiple manifestations from impaired social communication and language to restricted or repetitive behavior patterns, interests, and activities^1–3^. Due to the wide heterogeneity in behavior, and as recommended in The Diagnostic and Statistical Manual of Mental Disorders (DSM–5), this condition is referred as autism spectrum disorder (ASD), in which the term “spectrum” emphasizes the variation in the type and severity of manifestations^4^. ASD is thought to result from complex interactions during development between genetic, cellular, circuit, epigenetic and environmental factors^5–9^. Several researchers have suggested that an excitation/inhibition (E/I) imbalance during development^10,11^ may be an important mechanism, yet precise factors driving the disease are not well understood. Therapeutic interventions aiming to restore the E/I balance in ASD are then a major challenge^12^.

With regard to neurobiology, alterations in different brain networks have been found, e.g. in frontal, default mode and salience networks^13–18^, as well as in the social network^19^ – encompassing primary motor cortex, fusiform, amygdala, cerebellum, insula, somatosensory and anterior cingulate cortex^13,20,21^. ASD is also heterogeneous in relation to network characteristics; less segregation and greater efficiency has been shown^22,23^, and the opposite too^24^ or a combination of both^25,26^. Furthermore, ASD neuroanatomical correlates are not static but undergo changes throughout development^27–29^, and the same seems to occur behaviorally in social functioning and communication^30^. Altogether, accumulated evidence has shown high heterogeneity within ASD in the participation of functional brain networks and behavioral manifestations, but also, in the longitudinal trajectories at the single subject level.

Moreover, in relation to neuroimaging studies, recent work has shown additional sources of heterogeneity due to variations in the diagnostic and inclusion criteria, and differences in the processing neuroimaging pipeline^31,32^. ASD is also a polygenic highly heterogeneous condition, with 1010 different genes being associated with ASD as of July 8th 2022 according to the SFARI gene human-database, see also^33^. Of those, 213 have a relevance score of 1, meaning having maximum pathophysiological *published* evidence in relation to ASD. This high genetic complexity is also another manifestation of the heterogeneity in this condition. Previous work has assessed the relations between transcriptomics and brain morphology^34^, showing that genes which are down-regulated and enriched for synaptic transmission in individuals with autism were associated with variations in cortical thickness.

Novel strategies for ASD subtyping are needed to overcome such multi-scale heterogeneity, which is the largest drawback for the efficacy of therapies. Here, and following previous work^35–37^, we looked at large-scale brain connectivity patterns which are common within groups of patients to deploy subtyping in ASD. In particular, we applied consensus clustering strategies to multivariate connectivity patterns of brain regions^38,39^, and as a result, if two subjects belong to the same subtype, it means that each region in the two brains connects to other brain regions in a similar manner, revealing similarity in the multivariate brain connectivity of patients within the same subtype. Next, we linked connectivity-based ASD subtypes to their neurogenetic profile and hypothesized the ability to determine the biological processes that characterize each subtype, which is a major challenge in this condition. For this, we used the Allen Human Brain Atlas (AHBA) of whole-brain transcriptional data^40^ and characterized the functional connectivity patterns for each ASD subtype separately, following a strategy similar to the one used in several previous studies^41–47^. We performed our subtyping analyses on 657 autistic patients from the Autism Brain Imaging Data Exchange (ABIDE) repository^48^, all of them having passed a very strict quality-assurance criterion of elimination of subjects by head movement during image acquisition, thus correcting a well-known *spurious* excess of functional connectivity driven by head movements, which is even more pronounced in the autistic condition. Moreover, to overcome inter-scanner variability in the functional connectivity values across different Institutions, we applied rigorous harmonization strategies to transform data that are heterogeneous --and that come from different Institutions-- into equivalents^49–52^.

## Materials and Methods

### Participants

A total of N=2156 subjects from the ABIDE I^48^ and ABIDE II^53^ repositories were initially considered in this study, of which 1026 were ASD patients and 1130 were typically developing control (TDC) subjects. These data were collected across 35 different scanning Institutions. For each participant both anatomical and functional MRI data were used. Acquisition parameters for each scanning site are found at http://fcon_1000.projects.nitrc.org/indi/abide/. In addition, we also used the cognitive performance and disease severity information from the Autism Diagnostic Observation Schedule-Generic (ADOS-G), Autism Diagnostic Interview-Revised (ADI-R) and the verbal, performance and full Intelligence Quotient (IQ) scores (respectively, VIQ, PIQ and FIQ).

### Data quality-assurance

We discarded participants having scanning duration shorter than 5 minutes after scrubbing, lacking of full brain coverage, and with an average framewise displacement greater than 0.3 mm^54^. Participants from scanning studies KUL sample 3 and NYU sample 2 were also omitted because they only contained ASD subjects and therefore those cohorts did not provide any TDC. As a consequence, the number of finally considered subjects was 1541 (884 TDC, 657 ASD), corresponding to 33 scanning studies that were further merged into 24 institutions following the guidelines provided in http://fcon_1000.projects.nitrc.org/indi/abide/. The descriptive statistics per institution (number of subjects, ASD cases, mean age, sex distribution) is found in Table S1.

### Neuroimaging pre-processing and functional connectivity matrices

A state-of-the-art pre-processing pipeline was adopted using FSL 5.0.9, AFNI 16.0.01^55^ and MATLAB 2020b. We first applied slice-time correction, volume alignment to the average one to correct for head motion artifacts, which was followed by intensity normalization. We next regressed out 24 motion parameters, as well as the average cerebrospinal fluid (CSF) and average white matter signal. A band-pass filter was applied between 0.01 and 0.08 Hz, and linear and quadratic trends were removed. All voxels were spatially smoothed with a 6 mm FWHM. FreeSurfer v5.3.0 was used for brain segmentation and cortical parcellation. A total of 82 regions were generated, with 68 cortical regions from the Desikan-Killiany Atlas (34 in each hemisphere) and 14 subcortical regions (left/right thalamus, caudate, putamen, pallidum, hippocampus, amygdala, accumbens). The parcellation for each subject was projected to the individual functional data and the mean functional time series of each region was obtained. Finally, one connectivity matrix for each subject was built by Fisher z-transforming the Pearson correlation coefficients between the region pairs of time series. For structural neuroimaging group comparisons, region-wise volume and thickness were calculated with Freesurfer.

### Data Harmonization

To harmonize our multi-institution functional connectivity data, and before performing subtyping, we used an inhouse implementation of Combat (https://pypi.org/project/pycombat), adjusting batch effects by linear mixed modelling and the use of Empirical Bayes methods^50^. See suppl. information for details.

### ASD subtyping via consensus clustering

Consensus clustering was applied to brain connectivity matrices^38,39^. Since connectivity matrices may contain effects of not interest (e.g. age), prior to subtyping we regressed out age, sex and motion from each connectivity entry of the ASD subjects. To note, this regression-out step was only applied at this subtyping stage. In subsequent analyses, the original connectivity matrices were used and the effect of these variables was controlled for using them as covariates.

### Association between subtypes and transcriptomics

We computed the association between pseudo-R^2^ maps (more info at Suppl. Info) and brain transcriptomics maps using spatial autoregressive models to reduce the correlation-bias produced by the similar transcriptomic expression in proximal brain regions^45^. This analysis was implemented by means of the maximum-likelihood estimator routine (ML_Lag) from the Python Spatial Analysis Library (pysal)^56^. As a result, for each brain gene we obtained one t-stat and one p-value, which allowed us to assess the association with the pseudo-R^2^ maps while accounting for possible spatial autocorrelations. Among the significant associated genes, we identified as relevant those genes included in the SFARI database (https://gene.sfari.org/) with gene score equal to one.

### Gene set enrichment analysis and protein interaction analysis

We only considered for the analyses such genes with a p-value FDR corrected < 0.05 in each subtype. After that, we performed a Gene Set Enrichment Analysis (GSEA) using WebGestalt^57^ (http://www.webgestalt.org/) introducing as the input the list of the corrected genes and the t-stat from the association analysis, so the spatial correlation-bias was accounted for. We computed the GSEA for GO biological process^58^ and Reactome pathways^59^ and we only considered such enriched categories with a p-value FDR corrected < 0.05. For the protein interaction analysis, we used the tool STRING v11.5^60^ for generating a physical protein-protein interaction network for each subtype, with Experiments and Databases as interaction sources. These networks were after analyzed using Cytoscape v3.9.0.

## Results

We obtained harmonized functional connectivity matrices from 657 ASD and 884 typically developing control (TDC) subjects (Fig. 1). For subtyping, we first removed any effect from age, sex and head motion in the brain connectivity matrices of the ASD group and then applied a consensus clustering. We thus found two main subtypes^1^: the first one with 348 subjects (52.97% of all ASD subjects), and the second with 284 subjects (43.23%). In addition to these two subtypes, which were at the highest order in a hierarchy that was broken down into smaller subtypes (Fig. S1), we also found two residual subtypes of only 23 subjects (3.5%) and 2 subjects (0.3%) respectively, but they were ignored for further analysis due to their low number of subjects. The robustness of the subtyping solution was assessed by two different strategies, multi-resolution hierarchical clustering and cross-validation (suppl. info). As expected, given that their effects were removed prior to subtyping, none of the resulting subtypes were differentiated by age (absolute Cohen’s |d| = 0.04, t-test, p = 0.58), sex (Cramer’s V = 0.02, χ^2^ test, p = 0.55) or head motion (absolute Cohen’s |d| = 0.04, t-test, p = 0.64).

**Fig. 1.**
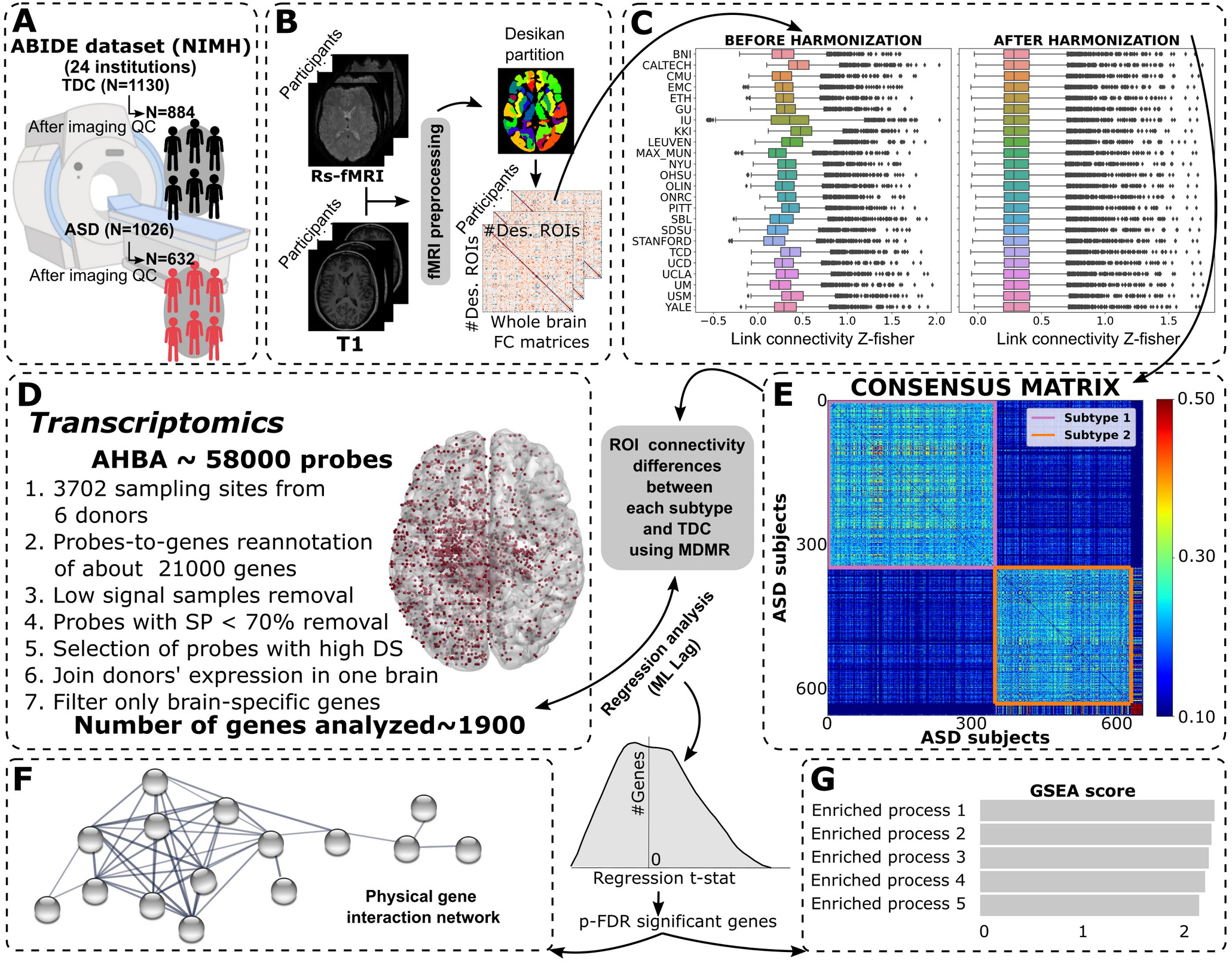
General workflow. **A:** Multicentric ABIDE dataset funded by NIMH and consisting in functional and anatomical MRI data from 24 different Institutions and two groups of subjects, TDC (N=1130) and ASD (N=1026). Some participants were eliminated after strict imaging quality checks (QC), resulting respectively in 884 TDC and 632 ASD. **B:** Image preprocessing and calculation of whole brain functional connectivity (FC) matrices using Desikan-Killiany *(Des.)* partition. **C:** Impact of rigorous data harmonization using the Combat algorithm to remove variability due to the effects of institution, age and sex. **D:** Transcriptome Allen Human Brain Atlas (AHBA) data preprocessing to be used for association with brain connectivity patterns in each subtype. **E:** ASD subtyping after consensus clustering applied to brain connectivity matrices. Using the results from D and E, we provide a biological characterization of each subtype using regression analysis between gene expression and connectivity patterns, and with a spatial autoregressive process for correlation-bias correction. **F:** Physical gene interaction network obtained only for the group of FDR-significant genes. **G:** Biological characterization of the significant associated genes using Gene Set Enrichment Analysis (GSEA).

With respect to cognitive and behavioral performance, none of the five scores provided significant differences (Table 1). Moreover, no significant differences between subtypes 1 and 2 were found on structural neuroimaging after multiple test corrections in either region volume or thickness. Therefore, all the following analyses are based on differences in functional connectivity that each ASD subtype has in in relation to TDC.

**Table 1:** Behavioral characterization of ASD subtypes. For each behavioral score, the number of observations in each subtype, their means and 95% confidence intervals, and the FDR-corrected p-values from a one-way ANOVA test to assess any statistical difference between them. Both ADI and ADOS total scores are composites of social and communication sub-item scores. For ABIDE data, we followed ADI_TOTAL = ADI_R_SOCIAL_TOTAL_A + ADI_R_VERBAL_TOTAL_BV, and ADOS_TOTAL = ADOS_COMM + ADOS_SOCIAL.

| SCORE<br>(N<sub>SUBTYPE 1</sub>/N<sub>SUBTYPE 2</sub>) | SUBTYPE 1 | SUBTYPE 2 | P-VALUE |
| --- | --- | --- | --- |
| **FIQ**<br>(321/266) | 105.23<br>[103.54-106.99] | 106.81<br>[104.96-108.61] | 0.31 |
| **VIQ**<br>(273/241) | 104.43<br>[102.28-106.51] | 106.78<br>[104.44-108.98] | 0.31 |
| **PIQ**<br>(277/245) | 105.0<br>[103.18- 106.96] | 105.4<br>[103.27- 107.56] | 0.80 |
| **ADI_TOTAL** | 35.21 | 34.07 | 0.31 |
| (228/190) | [34.07- 36.33] | [32.59- 35.51] |  |
| **ADOS_TOTAL** | 11.86 | 11.01 | 0.15 |
| (221/179) | [11.37- 12.36] | [10.46- 11.58] |  |

To assess the differences between groups in the overall connectivity per subject, defined here as the average positive correlation of the harmonized connectivity matrix, we performed an ordinary least squares (OLS) regression while controlling for age, sex and full intelligence quotient (FIQ) (Fig. 2). Subtype 1 showed significant hypo-connectivity with respect to TDC (*β* = −0.08, *p* < 0.01), and the opposite was true for the subtype 2 (*β* = 0.04, *p* <0.01), thus corresponding to hyper-connectivity. Moreover, the difference in (absolute) *β* coefficients, provided for subtype 1 higher values as compared to subtype 2, indicating a bigger separability in connectivity with respect to TDC. When using other metrics than the average positive correlation of the subject-level connectivity matrix, such as absolute value, median or trimmed mean, the same results were maintained (suppl. info).

**Fig. 2.**
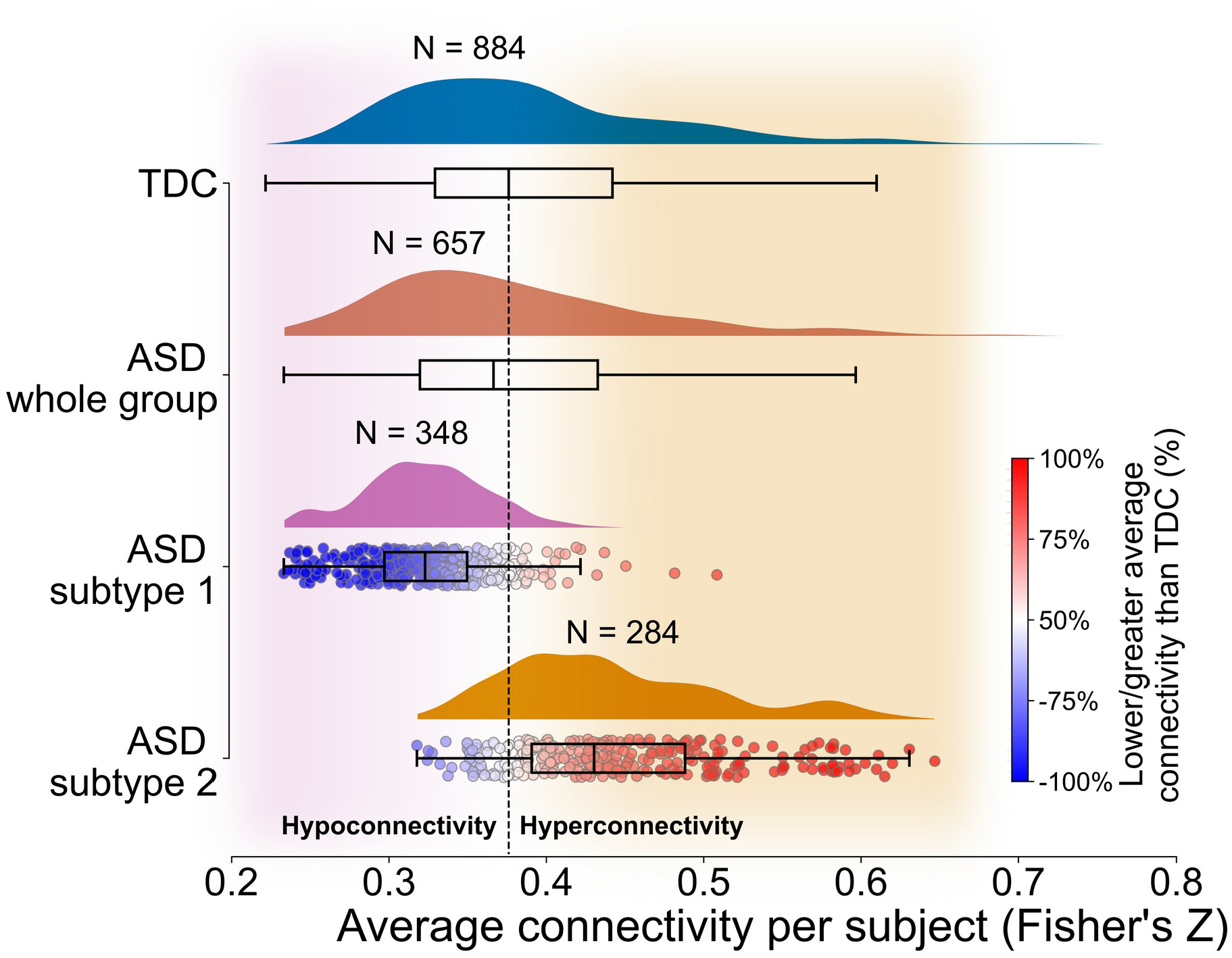
Two ASD subtypes, one with hypoconnectivity and the other with hyperconnectivity. Histogram and box plots of the individual average connectivity values (measured as Fisher’s Z) for the TDC group (blue), the population of all ASD subjects without (brown) and the two ASD subtypes (pink and orange). The median value of the TDC group is marked as the baseline by a dashed red line. Values greater than the baseline correspond to hyperconnectivity and those below the baseline hypoconnectivity. Subtype 1 is dominated by hypoconnectivity, and subtype 2 by hyperconnectivity. Additionally, within subtypes 1 and 2, we introduce two colors for the different subjects, blue for hypoconnectivity, and red for hyperconnectivity.

Next, we assessed the differences in connectivity pattern between each ASD subtype and the TDC group, measured by region-wise normalized pseudo-R^2^ statistical maps resulting from MDMR (Fig. 3, and Suppl. Inf.). The similarity between these spatial maps was very low (r=0.09, permutation-based p=0.67, after using 5000 surrogates that preserved spatial autocorrelation), indicating that each subtype exhibited a rather unique neurobiological profile of brain-wide connectivity patterns. Specifically, for subtype 1 higher differences as compared to TDC were found in superior temporal, posterior cingulate, and insula, covering the functional networks of default mode and salience. For subtype 2 higher differences existed in thalamus, putamen, and precentral, affecting to the networks of default mode and somatomotor. In this way, the two subtypes presented alterations in the connectivity patterns of regions within default mode network, but one subtype also showed specific alterations in salience (subtype 1) and the other in somatomotor (subtype 2).

**Fig. 3.**
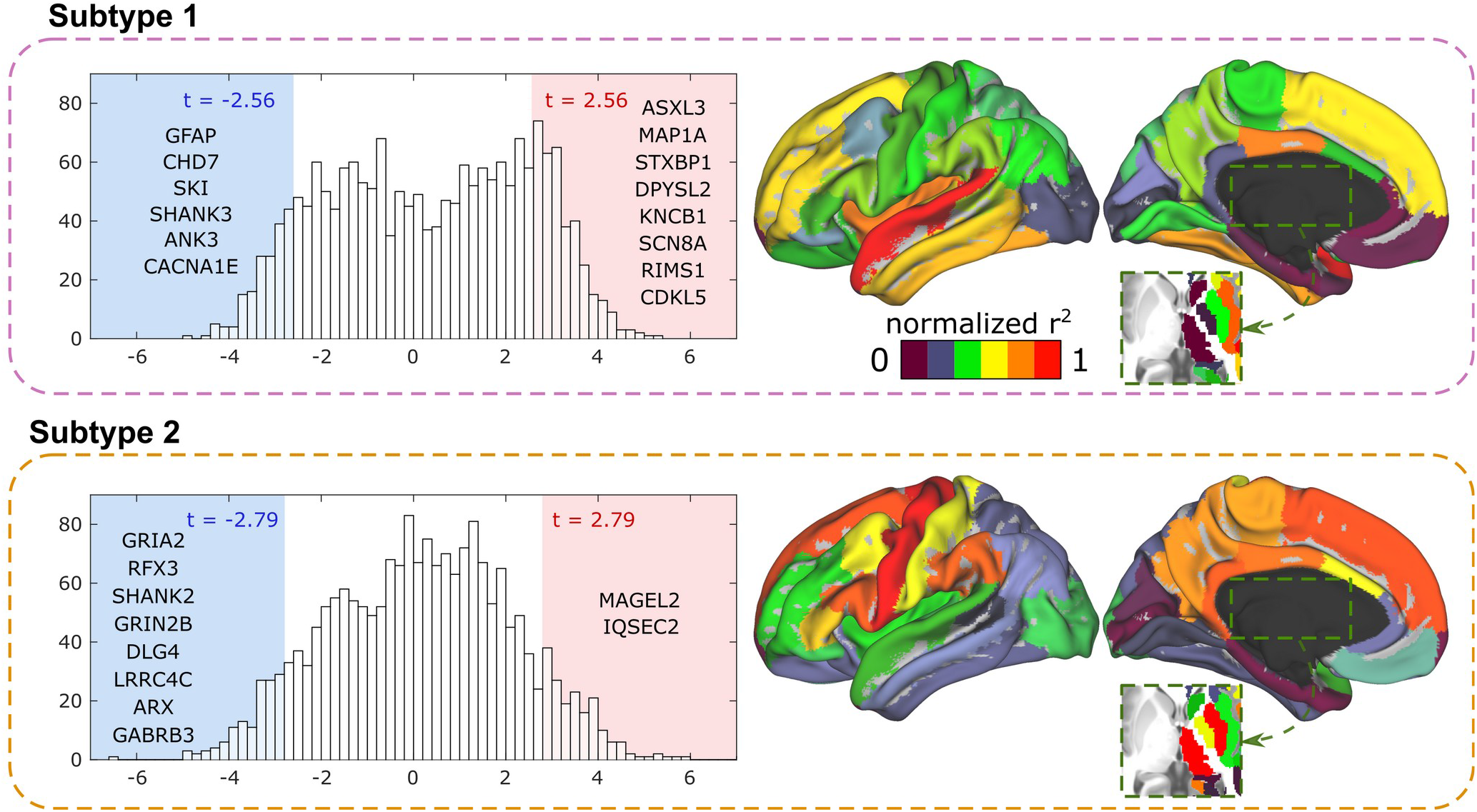
Association between transcriptome and connectivity patterns for each ASD subtype. For subtype 1 and 2, we calculated the pseudo-R^2^ vector (one component per brain region) accounting for the differences in connectivity pattern that each subtype has with respect to TDC. (Right) Brain maps of normalized pseudo-R^2^. (Left) Histograms of association values between pseudo-R^2^ and gene transcription activity (different values correspond to association with different genes). This procedure was repeated using the pseudo-R^2^ values for each subtype. The tail of the negative (N) genes (p-FDR < 0.05 and t-stat < 0) is marked by a blue rectangle and the tail of the positive (P) genes (p-FDR < 0.05 and t-stat > 0) by a red one for both subtypes. Significance-limits (t) are also shown. For each distribution tail, we also show the relevant genes present in the SFARI ASD genes with score = 1.

Subsequently, for the biological characterization of each subtype, we set out to identify which brain-related genes had an expression across brain regions significantly associated (p-FDR < 0.05) with the differences in connectivity measured by the normalized R^2^ brain maps (Fig. 3 - histograms). For subtype 1, a total of 195 negative-associated (NEG) genes and 364 positive-associated (POS) genes existed. Relevant NEG-genes were^2^ *GFAP, CHD7, SKI, SHANK3, ANK3*, and *CACNA1E*, while POS-genes were *ASXL3, MAP1A, STXBP1, DPYSL2, KNCB1, SCN8A, RIMS1*, and *CDKL5.* Similarly, for subtype 2, we found 142 NEG-genes, including *GRIA2, RFX3, SHANK2, GRIN2B, DLG4, LRRC4C, ARX, GABRB3*, and 180 POS-genes, including *MAGEL2* and *IQSEC2.* We next applied gene-enrichment to the list of significant genes within each subtype, finding no significant enrichment for subtype 1, the one with brain hypoconnectivity. However, for subtype 2, the enrichment of the NEG-genes included GO: Biological processes and Reactome pathways related to glutamate signaling (affecting both AMPA and NMDA receptors) and synapse organization, in relation with the excitation-inhibition imbalance occurring during development of brain circuits (Fig. 4A). We also assessed which NEG-genes participated in each biological process and pathway (Fig. 4B), finding that genes *DLG4, GRIN2B, GRIA2*, and *SHANK2* were participating in most of them; and in particular, the gene *DLG4* did it in all of them. In addition, the *DLG4* gene was the one with the highest degree in the protein interaction network.

**Fig. 4.**
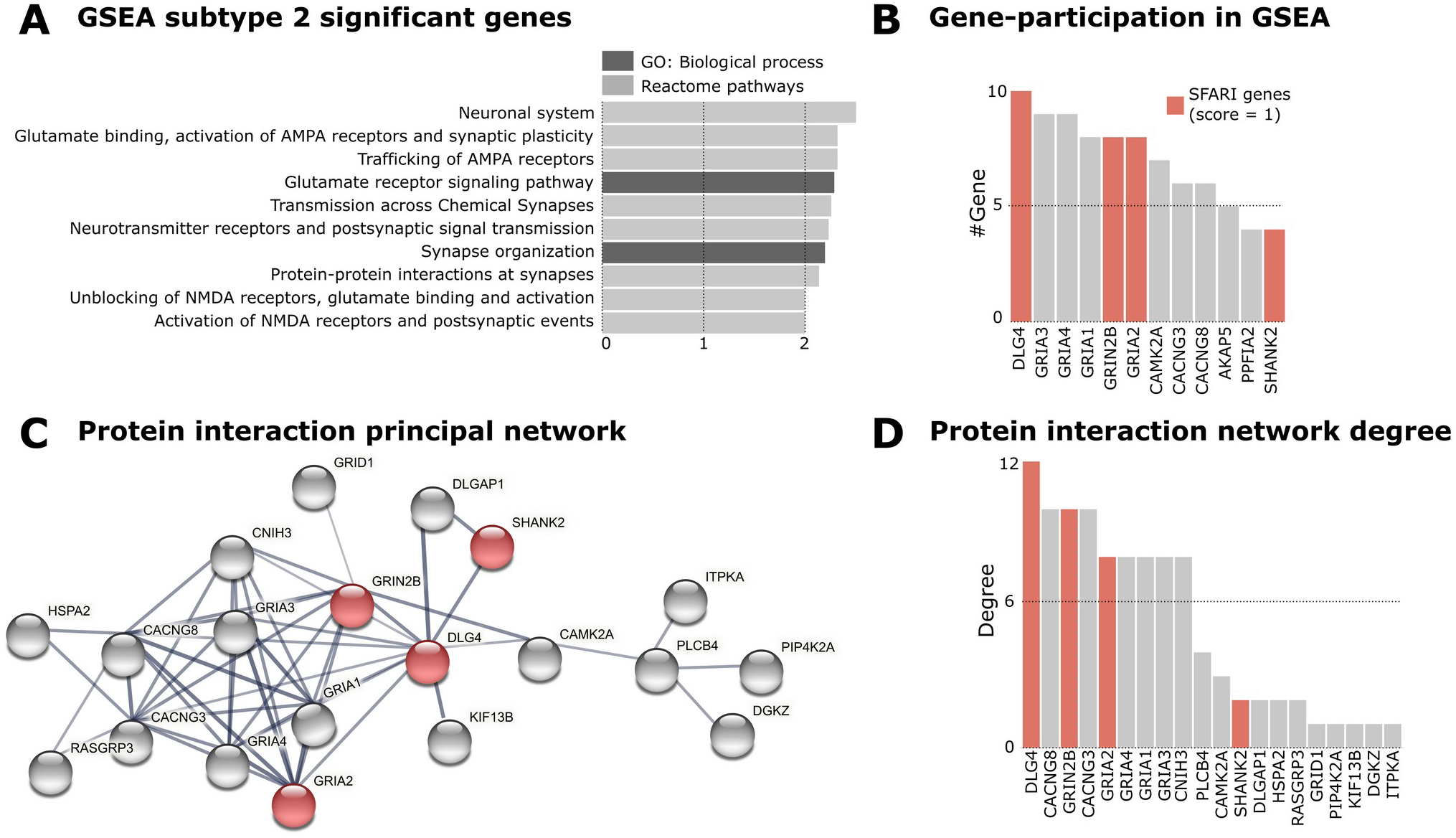
Excitation-Inhibition imbalance enrichment exists in one class of autistic subjects (Subtype 2). **A:** GSEA characterization of the FDR-significant genes in subtype 2, including the Gene Ontology (GO) Biological processes (dark gray) and Reactome pathways (light gray) enrichments. **B:** Participation count that each gene has in the processes shown in A, ranging from 4 to 10 (participating in all processes in panel A, only occurring for *DLG4).* **C:** Protein-protein interaction physical network from the list of FDR-significant genes. For the ease of visualization, only sub-networks with a minimum of 10 genes are depicted. **D:** Node degree of the genes participating in the network shown in C. *DLG4* is the gene with highest degree. **B,C:** Bars corresponding to genes with SFARI score =1 are colored in red, and the same occurs in D for network nodes.

Finally, it is important to note that no significant enrichment was found for subtypes lower in the dendrogram level corresponding to the two subtypes previously described (Fig. S1), neither after repeating the same analysis using the entire ASD group, thus indicating the need for subtyping to reveal our findings. Similarly, in order to test that the enrichment findings were inherent to the ASD group, we repeated our analytical pipeline over two matched subgroups of TDC subjects, one used for subtyping, and the other for estimating the pseudo-R^2^ maps with the resulting subtypes (see suppl. info). Two main subtypes of TDC emerged again, one characterized by hyperconnectivity and the other by hypoconnectivity. However, neither of these TDC subtypes provided genes that significantly associated with the pseudo-R^2^ vectors after correcting for multiple comparisons. Summing up, the significant association between excitation-inhibition imbalance and hyperconnected autistics is observed when subtyping in ASD and only in the ASD group, thus demonstrating the specificity of the reported enrichment.

## Discussion

Two major subtypes result from functional connectivity-based subtyping in a cohort of 657 autistic patients. The two are indistinguishable by morphometric comparisons based on structural neuroimaging and also with respect to behavioral scores. As compared to TDC, the first subtype is characterized by hypoconnectivity, connectivity alterations in default mode and salience networks with no significant gene enrichment after multiple comparisons. The second subtype, representing 43% of subjects with autism, is characterized by hyperconnectivity, network alterations in default mode and somatomotor networks with significant gene enrichment towards glutamate signaling (affecting both AMPA and NMDA receptors) and synapse organization, which is consistent with one of the most accepted hypotheses in the pathophysiology of autism, in relation to the excitation-inhibition imbalance which occurs during brain development. This enrichment is specific to the ASD condition and as such does not occur in the TDC group. Moreover, if no subtyping is performed, the connectivity profile in the entire autistic population has no significant enrichment, evidencing the need of *subtyping first* to find the connection towards excitation-inhibition imbalance in one class of autistic patients.

Some studies have set out to assess the heterogeneity in ASD for better stratifying the ASD condition^37,61–63^. Stratification yields reduced inter-individual differences and therefore, could complement -and even alleviate- the need for big sample sizes in autism-based biomarker discovery^64^. Our approach is unique in several ways. First, our study is based on a large cohort of patients with ASD (N = 657) from the ABIDE initiative, all of them having passed rigorous criterion of motion removal, and it combines anatomical and functional neuroimaging data from 24 different institutions. Second, we have used Combat, a rigorous data harmonization method to eliminate the variability between MRI scans across the 24 institutions, one of the largest sources of variability when combining imaging data from multiple institutions^65^. Third, our analysis of brain connectivity was carried out on a large-scale, where each brain region is represented by its connectivity pattern across the entire brain. Therefore, we do not consider *a priori* any brain region as more dominant or relevant than the others. Fourth, we made use of a consensus clustering approach we have developed^38,39^, and that has been successfully tested by others^66^, to group subjects in the same subtype if the connectivity vectors are similar across all the regions analyzed, which in our case involved a total of 68 cortical regions and 14 subcortical ones. Finally, we made use of the AHBA to describe the neurogenetic profiles of each subtype, which it has been used before for morphometric information in ASD^34^, but never for characterizing subtypes based on functional connectivity patterns of this condition.

Due to the large heterogeneity and diversity reported in ASD genetics, the use of AHBA may shed new light, as it provides information on the transcriptome across the brain in unprecedented detail, accounting for 3,702 sampling sites with transcription information of 20,500 genes as a specific signature for each anatomical region. Moreover, the use of AHBA is complementary to other techniques, such as genome-wide association studies or GWAS^67^, that simultaneously addresses genotype–phenotype associations from hundreds of thousands to millions of genetic variants in a data-driven manner. Indeed, GWAS has previously been used for ASD subtyping^68,69^, yet using behavioral scores as traits and therefore, the subtypes obtained were more closely related to symptom severity and not to functional connectivity.

Our results show that *DLG4*, a.k.a. *PSD95*, is the gene with major implication in the protein interaction network of subtype 2. *DLG4* mediates NMDA and AMPA receptor clustering and function, it affects glutamatergic transmission, and has been shown to have an aberrant function in ASD^70–74^. *DLG4* also influences the size and density of dendritic spines during brain development, having strong effects on synaptic connectivity and activity. In particular, reduced *DLG4* activity leads to increased dendritic spine numbers^75^, that might correspond to the hyperconnectivity found in these patients^10,76–78^ that in our case corresponded to those in subtype 2.

A number of limitations are present. First, our transcriptomic analysis made use of AHBA, which was obtained from brain samples of healthy donors and not from brain tissues of ASD patients, so the relations studied here between ASD-dependent connectivity patterns and healthy transcriptomics highlight large-scale organization aspects of the connectivity alterations in relation to gene expression. Future studies should confirm our findings using gene expression data from a pathologic cohort, which unfortunately is not currently available. A second limitation is that our study was restricted to the characterization of autism in terms of the functional networks that emerge from the resting activity of the brain, but we did not consider brain functionality across different tasks.

In summary, our novel approach, which includes data harmonization, multivariate distancing in large scale functional connectivity patterns and transcriptome brain maps, reveals strong enrichment in ASD for glutamate signaling (affecting both AMPA and NMDA receptors) and synapse organization, reinforcing the mechanistic hypothesis of excitation-inhibition imbalance occurring during development of autistic patients, linked to brain hyperconnectivity, but not to hypoconnectivity, thus suggesting a route for new potential therapeutic strategies in this subtype.

## Data and code availability

The data employed in this study belong to the ABIDE-I and ABIDE-II repositories. Their IDs can found in https://github.com/compneurobilbao/asd-subtyping-enrichment, as well as the codes used for the analyses.

## Acknowledgments

The authors are really pleased to thank Amaia Iribar Zabala for her fantastic work at the initial stage of this project and Dr. Rafael E. Oliveras-Rentas, Prof. Bennett Leventhal, Prof. Young Shin Kim, Prof. Oswald Quehenberger, Prof. Jose Delgado and Prof. Timothy Verstynen for the useful and insightful discussions. AJM is funded by a predoctoral contract from the Department of Education of the Basque Country (PRE-2019-1-0070). MTH and JMC are funded by Ikerbasque: The Basque Foundation for Science. JMC is funded by the Department of Economic Development and Infrastructure of the Basque Country (Elkartek Program Grant KK-2021-00009).

## Abbreviations

ABIDE: Autism Brain Imaging Data Exchange
ADI-R: Autism Diagnostic Interview-Revised
ADOS-G: Autism Diagnostic Observation Schedule-Generic
AHBA: Allen Human Brain Atlas
ASD: Autism Spectrum Disorder
CSF: cerebrospinal fluid
Des: Desikan-Killiany
DSM-5: The Diagnostic and Statistical Manual of Mental Disorders V
E/I: Excitation/Inhibition
FC: Functional Connectivity
FDR: False Discovery Rate
FIQ: Full Intelligence Quotient
GO: Gene Ontology
GSEA: Gene Set Enrichment Analysis
GWAS: Genome-Wide Association Studies
MDMR: Multivariate Distance Matrix Regression
MRI: Magnetic Resonance Imaging
NIMH: National Institute of Mental Health
NEG: Negative Associated
OLS: Ordinary Least Squares
PIQ: Performance Intelligence Quotient
POS: Positive Associated
QC: Quality Checks
SFARI: Simons Foundation Autism Research Initiative
TDC: Typically Developing Control
VIQ: Verbal Intelligence Quotient

## Supplementary Information

### Data Harmonization

Batch effects may reflect the different set-ups for image acquisition at each institution included (e.g., MRI scanner manufacturer, different antenna and/or software, gradient coils, magnet field strength, etc.). Let *Y_ijk_* represent the value of the connectivity entry *k* for subject *j* at institution *i.* Combat adjusts the *Y_ijk_* data by estimating the coefficients present in the following linear mixed model:

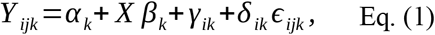

where *α_k_* is the fixed intercept, *β_k_* the fixed slopes for the variables in a design matrix *X*, and *γ_ik_* and *δ_ik_* the location and scale institution factors modelled as random effects. One of the strong points of Combat is the use of an empirical Bayes (EB) approach to better estimate *γ_ik_* and *δ_ik_*, an iterative step that is particularly relevant when sample sizes are small. Specifically, it assumes that the two parameters controlling the random effects are sampled from the following prior distribution:

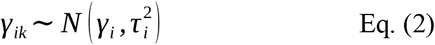

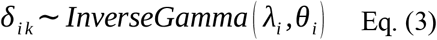

The hyperparameters *γ_i_*, 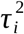, λ_*i*_ and *θ_i_* are empirically estimated using an expectation-maximisation (EM) procedure as described in^1^. Thus, the harmonized data 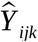 read:

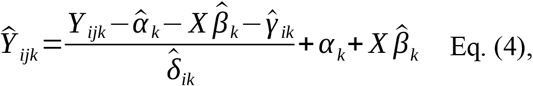

where 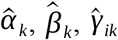 and 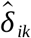 are the fitted coefficients present in Eq. (1). The design matrix *X* encodes the effects of interest that we want to preserve during the harmonization process, which in our case corresponded to the diagnosis label (TDC vs ASD) in connectivity.

We verified the presence of heterogeneity related to scanning institution in our connectivity matrices by applying to each link a Kruskal-Wallis test, which aims at testing median location differences, and a Lavene’s test, which assesses differences in variances. The former yielded all the links significantly different across institutions after FDR correction (3321), whereas the latter gave 1236. Such scanning institution differences disappeared after the application of Combat, while retaining the between-group variability of our data.

### ASD subtyping via consensus clustering

First, we regressed out from the ASD harmonized connectivity matrices the effects of age, sex and head motion, since they could affect subsequent subtyping. Then, these matrices were used to define, for each brain region *i*, a matrix of euclidean distances between *u* and *v*ASD subjects, i.e.

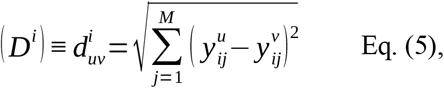

where *M* = 82 reflects the number of brain regions, and 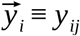 the whole-brain connectivity pattern for a given region *i*, i.e. a vector of dimension equal to the number of regions, where each component is defined as the amount of connectivity between the given region and any other in the atlas. Then, each distance matrix *D^i^* was partitioned into *k* groups of subjects using a k-medoids clustering method^2^, and the resulting clustering information encoded into an adjacency matrix, whose entries are 1 if a pair of subjects belongs to the same cluster and zero otherwise. Subsequently, a *N* × *N* consensus matrix *C* was evaluated by averaging this information across the nodes. Hence, the entries of *C_uv_* indicate the number of partitions in which subjects *u* and *v* are assigned to the same group, divided by the number of partitions. Eventually, the consensus matrix is averaged over the *k* range in the interval (2-20), so that information about the underlying structure at different resolutions is combined in the final consensus matrix, for more details see^3^. The consensus matrix *C* was further used to define a Newman and Girvan-like modularity matrix^4^:

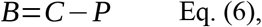

where *P* is the expected co-assignment matrix, uniform as a consequence of the null ensemble strategy obtained by repeating the permutation of labels 1000 times. Such a modularity matrix *B* encodes all the information about the interaction between subjects at different levels. As a result, one could now define any distance quantity applied to this matrix for assessing clustering. Instead, we directly fed this *B* matrix into a generalized Louvain method for community detection (https://github.com/GenLouvain/GenLouvain), yielding an optimal output partition that maximizes the network modularity.

### Robustness of subtyping solution through multi-resolution hierarchical clustering

Given the implicit resolution dependence when clustering of the Newman and Girvan’s modularity matrix^5^, we validated our subtyping solution by applying to the consensus matrix a multi-resolution clustering method that finds the hierarchical community structure after sampling the entire ranges of possible resolutions^6^. As a result of this, two major clusters were obtained, which remain fairly stable and invariant as we go down the multi-resolution hierarchical tree. When comparing these major clusters in the higher level of the tree with our two original subtypes, we found a large similarity (Normalized Mutual Information NMI = 0.91) and no statistical differences in the subjects content (χ^2^=0.15, p = 0.69) between both clustering solutions.

### Robustness of subtyping solution through cross-validation

We first split the raw data into two halves matched by age, sex and head motion and then the same harmonization and clustering procedure was applied to both subsamples. The first half yielded two ASD subtypes (189 and 139 subjects), whereas the second half also produced two subtypes of comparable subjects (179 and 142 subjects), in addition to two residual subtypes (7 and 1 subjects). Subsequently, a connectome-based predictive modeling^7^ was trained on the first half (observed labels provided by its subtyping solution) and then tested on the observations of the two major subtypes in the second half. The accuracy between predicted and observed labels (i.e. those provided by its subtyping solution) was 93.41%, showing a high agreement between both subsamples with respect to the presence of two similar ASD subtypes, and thus confirming the reliability of our subtyping results.

### Separability of ASD subtypes with respect to TDC

To assess the separability of brain connectivity profiles between each ASD subtype and the TDC group, we performed a *Multivariate Distance Matrix Regression* (MDMR) analysis^8,9^. Specifically, MDMR regressed each distance matrix per region given by Eq. (5) onto a design matrix *X* formed by a set of *m* predictors, yielding a pseudo-F statistic 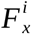 that reads:

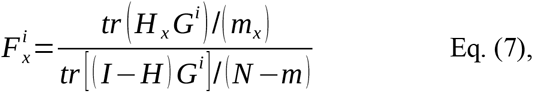

where *tr* indicates the trace operator, *N* the number of observations, *m_x_* the degrees of freedom of predictor *x, H_x_* the isolated effect of predictor *x* from the usual matrix *H* = *X*(*X^T^ X*)^-1^ *X^T^*, and *G^i^* the so-called Gower matrix built from the distance matrix *D^i^*^10^. In our case, the predictors consisted of the ASD vs TDC group factor as the variable of interest, and sex, age, mean framewise displacement and FIQ as covariates. Like the F-estimator in a standard ANOVA analysis, Eq. (3) assesses the variance explained by a predictor variable with respect to the unexplained variance. Finally, to estimate how much variability can be attributed to each predictor, a pseudo-R^2^ effect size can be computed by dividing the numerator without the degrees of freedom in Eq. (3) by the total sum of squared pairwise distances in the Gower matrix

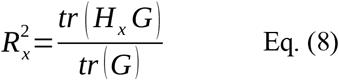

Similar to standard linear models, this effect size quantifies the proportion of the total sum of squares that can be explained by the predictors.

### Robustness of hyper- and hypo-connectivity findings

To assess the connectivity class in each subtype, we compared the average connectivity per subject between each subtype and the TDC group (baseline). We found hypo-connectivity for subtype 1 and hyper-connectivity for subtype 2, both situations with respect to the TDC group (baseline). These results were obtained when comparing the positive values of the correlation matrix, but the same results were found when using the absolute correlation matrix, or the median or trimmed means as the overall connectivity metric, preserving in all cases the findings of hypo-connectivity for subtype 1 and hyper-connectivity for subtype 2.

### Transcriptomics

To build brain transcription maps, we took advantage of the publicly available data in the AHBA^11^. The dataset consisted of MRI images, and a total of 58,692 microarray-based transcription profiles of about 20,945 genes sampled from 3,702 different regions across the brains of six humans. To pool all the transcription data into a single brain template, we followed a similar procedure to that employed elsewhere^12^, which includes: (1) Probe reannotation using a re-annotator toolkit^13^; (2) Removal of probes whose sampling proportion in any of the six brains did not exceed the 70%; (3) Unique probe to gene assignment using the maximum differential stability (DS) criterion^14^; (4) Removal of the inter-subject differences by pooling together the Z-scores of the transcription values for each gene and brain; (5) Computation of a single transcription value for each region in the left hemisphere of the Desikan-Killiany atlas by calculating the median of all the values belonging to the given region; (6) Gene filtering considering only brain-specific genes relative to other tissues using the Human Protein Atlas^15,16^ (https://www.proteinatlas.org).

### Robustness of association between ASD subtypes and transcriptomics

We also tested if the significant genes resulting from association between pseudo-R^2^ statistical maps and gene expression were dependent on the clustering method used, comparing the results of generalized Louvain algorithm to those found by multiresolution clustering. As a measure of distance between the two solutions, we computed the dice index between the gene list associated with subtype 1 and the same for subtype 2, obtaining respectively dice values of 0.99 and 0.89, which indicating high-level of reproducibility of the gene-expression association to brain alterations between the two clustering strategies.

Finally, to prove that our gene and enrichment findings were exclusive of the ASD condition, we also repeated the same procedure but only in the TDC population. To do that, we first divided the entire TDC cohort into two subgroups, half sized and randomly chosen so that they were well-matched for age, sex, movement, FIQ, and overall connectivity (mean across positive entries in the connectivity matrices). Next, we applied in one subgroup the same subtyping procedure (including the same previous denoising step to avoid clusters driven by effects of not interest), obtaining again two subtypes, one representing hyperconnectivity and the other hypoconnectivity. We then used the other TDC subgroup to calculate the pseudo-R^2^ statistical maps, which were subsequently associated with the transcriptomics data. As a result, no gene survived by FDR in any subtype, thus concluding that the excitation-inhibition imbalance found in the hyperconnected autistic subtype is specific to the autistic condition.

---

**Table S1:** Main data characteristics for each Institution participating in our study. BNI = Barrow Neurological Institute; CALTECH = California Institute of Technology; CMU = Carnegie Mellon University; EMC = Erasmus University Medical Center Rotterdam; ETH = ETH Zürich; GU = Georgetown University; IU = Indiana University; KKI = Kennedy Krieger Institute; LEUVEN = University of Leuven; MAX_MUN = Ludwig Maximilians University Munich; NYU = NYU Langone Medical Center; OHSU = Oregon Health and Science University; OLIN = Olin; Institute of Living at Hartford Hospital; ONRC = Olin Neuropsychiatry Research Center, Institute of Living at Hartford Hospital; PITT = University of Pittsburgh School of Medicine; SBL = Social Brain Lab, Netherlands Institute for Neurosciences; SDSU = San Diego State University; STANFORD = Stanford University; TCD = Trinity Centre for Health Sciences; UCD = University of California Davis; UCLA = University of California Los Angeles; UM = University of Michigan; USM = University of Utah School of Medicine; YALE = Yale Child Study Center. I: ABIDE 1. II: ABIDE 2.

| ***Institution name*** | ***Number of***<br>***subjects***<br>***contributed*** | ***Age***<br>***(mean ± sd)*** | ***Sex***<br>***distribution***<br>***(Female)*** | ***Number of***<br>***ASD cases*** |
| --- | --- | --- | --- | --- |
| ***BNI (II)*** | 40 | 39.85 ± 15.59 | 0 | 19 |
| ***CALTECH (I)*** | 37 | 27.42 ± 9.76 | 8 | 18 |
| ***CMU (I)*** | 20 | 25.45 ± 5.29 | 4 | 8 |
| ***EMC (II)*** | 27 | 8.40 ± 1.09 | 4 | 15 |
| ***ETH (II)*** | 26 | 22.91 ± 4.57 | 0 | 6 |
| ***GU (II)*** | 76 | 10.92 ± 1.66 | 28 | 29 |
| ***IU (II)*** | 37 | 24.43 ± 7.59 | 9 | 17 |
| ***KKI (I and II)*** | 205 | 10.29 ± 1.28 | 70 | 48 |
| ***LEUVEN (I)*** | 61 | 18.18 ± 4.97 | 7 | 26 |
| ***MAX_MUN (I)*** | 44 | 28.77 ± 11.79 | 7 | 18 |
| ***NYU (I and II)*** | 245 | 13.82 ± 6.74 | 42 | 115 |
| ***OHSU (II)*** | 83 | 10.95 ± 2.05 | 31 | 33 |
| ***OLIN (I)*** | 22 | 17.41 ± 3.70 | 4 | 12 |
| ***ONRC (II)*** | 18 | 22.39 ± 3.58 | 5 | 7 |
| ***PITT (I)*** | 43 | 19.53 ± 6.87 | 7 | 22 |
| ***SBL (I)*** | 25 | 34.08 ± 6.41 | 0 | 13 |
| ***SDSU (I and II)*** | 86 | 13.75 ± 2.67 | 15 | 42 |
| ***STANFORD (I)*** | 19 | 10.23 ± 1.48 | 6 | 11 |
| ***TCD (I and II)*** | 73 | 16.86 ± 3.41 | 0 | 33 |
| ***UCD (II)*** | 28 | 14.98 ± 1.81 | 7 | 15 |
| ***UCLA (I and II)*** | 70 | 12.98 ± 2.42 | 8 | 36 |
| ***UM (I)*** | 107 | 14.60 ± 3.26 | 24 | 41 |
| ***USM (I and II)*** | 103 | 23.36 ± 7.67 | 5 | 52 |
| ***YALE (I)*** | 46 | 12.97 ± 2.97 | 13 | 21 |
| ***ALL*** | 1541 | 16.50 ± 8.82 | 304 | 657 |

**Figure S1:**
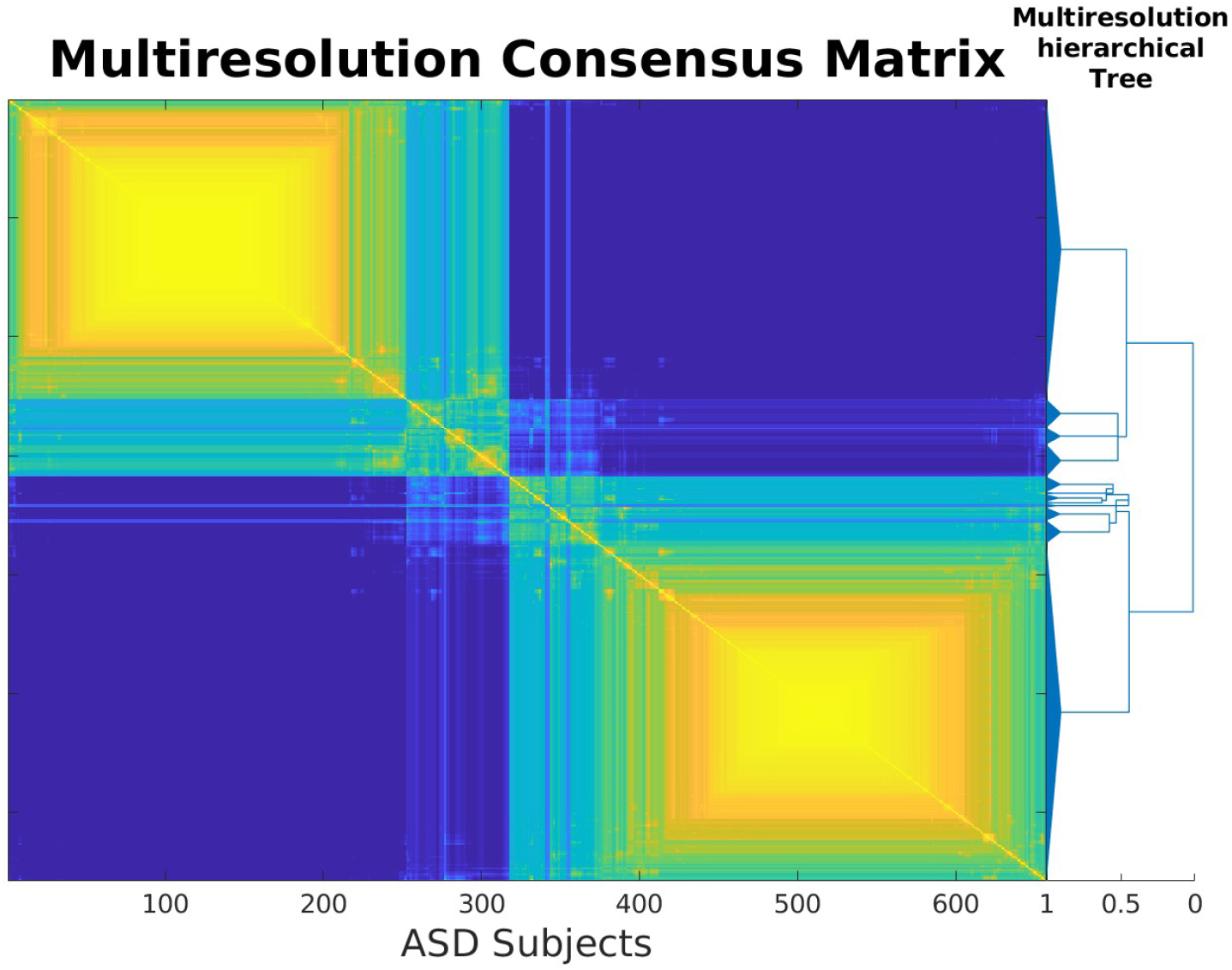
The two major ASD subtypes were subdivided hierarchically into smaller subtypes, but the significant enrichment found only occurred for the two subtypes at the highest dendrogram level.

## Footnotes

1 For subtyping, we only use connectivity matrices from subjects with ASD. The matrices from TDC subjects were only used for the neurobiological characterization of each subtype, as we studied the association between the separability of each ASD subtype with respect to TDC, encoded in the pseudo-R2 vectors, and brain transcriptomics.

2 Genes with a significant association (p-FDR < 0.05) and present in the SFARI gene human-database with a relevance score of 1.

